# Multimodal brain features at preschool age and the relationship with pre-reading measures one year later: an exploratory study

**DOI:** 10.1101/2021.04.30.442198

**Authors:** Kathryn Y. Manning, Jess E. Reynolds, Xiangyu Long, Alberto Llera, Deborah Dewey, Catherine Lebel

## Abstract

Pre-reading language skills develop rapidly in early childhood and are related to brain structure and function in young children prior to formal education. However, the early neurobiological development that support these skills is not well understood and has not been assessed longitudinally using multiple imaging approaches. Here we acquired anatomical, diffusion tensor imaging (DTI) and resting state functional MRI (rs-fMRI) from 35 children at 3.5 years of age. Children were assessed for pre-reading abilities using the NEPSY-II subtests one year later (4.5 years). We applied a data-driven linked independent component analysis to explore the shared co-variation of grey and white matter measures. Two sources of structural variation at 3.5 years of age demonstrated a weak relationship with Speeded Naming scores at 4.5 years of age. The first imaging component involved volumetric variability in reading-related cortical regions alongside microstructural features of the superior longitudinal fasciculus. The second component was dominated by cortical volumetric variations within the cerebellum and visual association area. In a subset of children with rs-fMRI data, we evaluated the inter-network functional connectivity of the left-lateralized fronto-parietal language (FPL) network and its relationship with pre-reading measures. Higher functional connectivity between the FPL functional network and the default mode and visual networks at 3.5 years predicted better Phonological Processing scores at 4.5 years. Together, these results suggest that the integration of functional networks, as well as the co-development of white and grey matter brain structures in early childhood, may support the emergence of pre-reading measures in preschool children.

## Introduction

Reading is an essential skill that plays a fundamental role in academic achievement, social engagement with the world and peers, and mental health (Dehaene-Lambertz et al., 2006; Torppa et al., 2006; May et al., 2011; Lonigan et al., 2013). Language skills develop dramatically throughout early childhood and lay the foundation for the reading skills children will develop. Phonological awareness and speeded naming skills in young children are strong predictors of future reading success (Lonigan et al., 2009; Snowling and Hulme, 2011). The neurobiological processes that support development of these skills are primed in utero (May et al., 2011) and develop rapidly during early life. A thorough understanding of the regionally-specific structural and functional brain features associated with later reading abilities is essential to determine how reading difficulties may be related to early brain differences, and to help appropriately target early language and reading interventions (Zijlstra et al., 2020).

Reading involves a left lateralized network of both grey matter regions and white matter connections in the brain (Torppa et al., 2006; Zijlstra et al., 2020). Frontal and temporal-parietal regions are connected via dorsal white matter pathways (arcuate fasciculus, superior longitudinal fasciculus [SLF]), and frontal, temporal, and occipital regions are connected by ventral white matter pathways (inferior longitudinal [ILF], inferior fronto-occipital [IFO], and uncinate fasciculi [UF]) (Klingberg et al., 2000; Brown et al., 2001; Welcome et al., 2011). Left lateralization is observed in both functional brain connectivity patterns (Benischek et al., 2020) and structurally (Lebel et al., 2013), with increased lateralization linked to language skill development (Reynolds et al., 2019b).

Relationships between reading performance and brain structure have been observed in adults (Klingberg et al., 2000; Brown et al., 2001; Welcome et al., 2011), adolescents (Kronbichler et al., 2008; Lebel et al., 2013), and school-aged children that are just beginning to read (Beaulieu et al., 2005; Deutsch et al., 2005; Eckert et al., 2005; Niogi and McCandliss, 2006; Lebel and Beaulieu, 2009). Children aged 6-7 years who subsequently received a diagnosis of dyslexia had reduced cortical thickness in left hemisphere reading-related regions compared to children who did not eventually receive a diagnosis (Clark et al., 2014). While neuroimaging studies have identified isolated imaging measures associated with reading ability in school-age children, the neurobiological development underlying the development of reading skills may begin earlier in life (Zuk et al., 2019). Even before formal reading education, the development of language and pre-reading skills are supported by the developing structural and functional architecture of the brain. Recent research exploring brain functional architecture in pre-readers during early childhood demonstrates relationships between reading-related gray and white matter maturation and pre-reading skills (Raschle et al., 2012; Saygin et al., 2013; Walton et al., 2018; Reynolds et al., 2019b) suggesting that differences in brain functional connectivity may influence later reading acquisition. A recent cross-sectional passive viewing fMRI study (Benischek et al., 2020) identified stronger connectivity between reading regions, increased connectivity between reading and motor areas, and decreased connectivity with default mode network and visual regions in children who had higher pre-reading skills. Furthermore, pre-readers with a family history of dyslexia show reduced temporal-parietal and temporal-occipital activation during sound matching compared to pre-readers without a family history of dyslexia (Raschle et al., 2012).

Longitudinal studies have shown that faster changes in fractional anisotropy (FA) and mean diffusivity (MD) are associated with larger gains in reading skills in children with and without developmental disorders (Yeatman et al., 2012a; Treit et al., 2013; Wang et al., 2017). Cortical structure in children and adolescents has been associated with phonological processing skills (Lu et al., 2007), with higher baseline reading skills associated with faster changes in gray matter volume in typically developing children aged 5 to 15 years (Linkersdörfer et al., 2014). Furthermore, changes in microstructure have been demonstrated following intensive reading interventions (Keller and Just, 2009; Huber et al., 2018). Importantly, longitudinal studies indicate activation and functional connectivity in reading related regions are related to language skill gains (Xiao et al., 2016) and later reading outcomes (Jasińska et al., 2020) during early childhood. In general, children with higher reading proficiency show faster development of reading-related brain regions than children with poor reading skills (Yeatman et al., 2012b; Wang et al., 2017; Reynolds et al., 2019b). While there are few longitudinal studies focused on reading and language development during the pre-school period, one recent study reports that infant arcuate fasciculus and corticospinal tract microstructure predicts phonological processing and vocabulary at 5 years of age (Zuk et al., 2019). These studies suggest associations between early brain structure or function and the development of reading skills in children, but how multimodal neurobiological features underlie pre-reading measures in preschool aged children, before formal reading education, is still not well-understood.

Neurodevelopment involves concurrent changes in both white and gray matter structures as well as brain functional communication and architecture. An understanding of the multi-faceted imaging measures that underlie language in early childhood may help improve our understanding of the neurobiological development that supports reading later in life. While brain structure, microstructure and function are often investigated in isolation, they are inherently linked to one another throughout development. Acquiring and analyzing multiple imaging sequences simultaneously can provide a more thorough interpretation of the complex neurobiology that underlies changes in these measures. In this study, we applied a data-driven linked independent component analysis technique (Llera et al., 2019) to explore structural and microstructural co-variations that relate to pre-reading measures. This approach has been used to study healthy development and aging as well as childhood disorders like autism spectrum disorder and ADHD (Itahashi et al., 2015; Wolfers et al., 2017). The aim of this longitudinal study was to explore early childhood brain morphometry, white matter microstructure, and functional resting state networks in conjunction to identify regionally specific neurobiological features that may predict measures of pre-reading skills later in childhood.

## Methods

### Participants

Participants were 35 young children (19 boys/16 girls) selected from the larger ongoing Calgary Preschool MRI Study (Reynolds et al., 2020) based on the criteria of having high-quality T1-weighted and diffusion weighted scans at 3.5 years (3.49 ± 0.14; range 3.25-3.75 years) and a pre-reading assessment 4.5 years (4.50 ± 0.16; range 4.25-4.75 years). The University of Calgary Conjoint Health Research Ethics Board (CHREB) approved this study (REB13-0020). Informed written consent was obtained from each participant’s legal guardian prior to the commencement of the study, and ongoing verbal assent was obtained from the participants.

### Data Availability

Imaging data from the Calgary Preschool MRI study is available through an Open Science Framework (https://osf.io/axz5r/).

### Language assessments

Children had their pre-reading skills assessed using the NEPSY-II Speeded Naming and Phonological Processing subtests (approx 20 mins) at 4.5 years of age. The Speeded Naming subtest assesses rapid semantic access to and production of names of colors and shapes, and the Phonological Processing subtest assesses phonemic awareness (Korkman et al., 2007). Age standardised Speeded Naming Combined Scaled Scores (accounts for both speed and accuracy) and Phonological Processing Scaled Scores were calculated and used in the analysis.

### MRI acquisition

All imaging was conducted using the same General Electric 3T MR750w system and a 32-channel head coil (GE, Waukesha, WI) at the Alberta Children’s Hospital in Calgary, Canada. Children were scanned either while awake and watching a movie of their choice, or while sleeping without sedation. fMRI scans during which the child was asleep were excluded from analyses. Prior to scanning, parents were provided with detailed information on MRI procedures and given the option to complete a practice MRI session in a training scanner to familiarize the child with the scanning environment, or to make use of a take home pack with this information (e.g., noise recordings (Thieba et al., 2018)). Families were also provided with a book that incorporates our scanning procedures into an engaging story (Frayne, 2015) and we encouraged the parents/guardians to review the materials with the child.

T1-weighted images were acquired using a FSPGR BRAVO sequence, 210 axial slices; 0.9 x 0.9 x 0.9mm resolution, TR = 8.23 ms, TE = 3.76 ms, flip angle = 12°, matrix size = 512 x 512, inversion time = 540 ms. Whole-brain diffusion weighted images were acquired using single shot spin echo echo-planar imaging sequence: 1.6 x 1.6 x 2.2 mm resolution (resampled on scanner to 0.78 x 0.78 x 2.2 mm), full brain coverage, TR = 6750 ms; TE = 79 ms (set to minimum for first year), 30 gradient encoding directions at b=750s/mm^2^, and five interleaved images without gradient encoding at b=0s/mm^2^ for a total acquisition time of approximately four minutes. Passive viewing fMRI data were acquired with a gradient-echo echo-planar imaging (EPI) sequence, 36 axial slices, 3.59 x 3.59 x 3.6 mm resolution, TR = 2000 ms, TE = 30 ms, flip angle = 60°, matrix size = 64 x 64, 250 volumes.

### Image processing

#### T1-Weighted

Voxel-based morphometry (VBM) processing was undertaken on the T1-weighted images using FSL to create gray matter maps. N4-bias correction was performed using ANTs (Avants et al., 2014), then images were transformed to radiological orientation. Brain extraction was undertaken using BET via the standard FSL VBM protocol; when brain extraction using the default settings was unsuccessful (n=7), BET was performed manually to achieve the best results. Next, images were segmented into gray matter, white matter and CSF. The gray matter images were then affine registered to the NIHPD asymmetrical pediatric brain template (4.5 to 8.5 years template; this template was used to be consistent with our prior work in the larger Calgary Preschool MRI sample and to permit future analysis spanning 2-8 years) in Montreal Neurological Institute (MNI) standard space (Fonov et al., 2011), resampled to 2mm isotropic voxels, concatenated and averaged. Using the standard FSL pipeline, these average gray matter images were used to create a study-specific non-linear gray matter template (2mm isotropic voxels), following which gray matter images were non-linearly registered to the study-specific template, modulated, smoothed (2mm), and concatenated into one 4D file.

#### Diffusion MRI

Raw diffusion images were visually quality checked and all motion-corrupted volumes, or volumes with artifacts were removed prior to processing. Datasets passed final quality assurance checks (n = 35) if they had at least 18 high quality diffusion weighted volumes, and two high quality b0 volumes remaining following volume removal. Data was then pipelined through ExploreDTI V4.8.6 (Leemans et al., 2009) to correct for signal drift, Gibbs ringing (non-DWIs), subject motion, and eddy current distortions. Fractional anisotropy (FA), axial- (AD), and radial diffusivity (RD) image maps were extracted. For images where brain extraction during DTI preprocessing did not remove all non-brain material, FSL BET was run on the extracted FA map, and the resulting binary mask was used to mask the FA, AD, and RD images. These DTI maps were then non-linearly warped using ANTs (Avants et al., 2014) to the NIHPD asymmetrical pediatric brain template (ages 4.5-8.5 years) in Montreal Neurological Institute (MNI) standard space (Fonov et al., 2011). All of the registered diffusion data was then merged into one four-dimensional image to create a mean fractional anisotropy (FA) mask and image for all subjects. The mean FA image was skeletonised with a threshold of FA > 0.2 to create a mean FA skeleton mask. All participants FA, AD and RD images were projected onto that skeleton.

Analysis was conducted both on these skeletonized whole-brain maps of diffusion measures, as well as on a subset of tracts that have been associated with reading in children: the bilateral uncinate fasciculus (UF), inferior longitudinal fasciculus (ILF), inferior fronto-occipital (IFO), and superior longitudinal fasciculus (SLF). Tracts were identified using the JHU white matter tractography atlas.

### Linked Independent Component Analysis

Gray matter morphometry, skeletonised FA, AD and RD maps were aligned to the same space, concatenated, and then used as input for a linked independent component analysis using FSL tools (Llera et al., 2019). This is a data-driven approach that uncovers the neurobiological variations across multiple modalities or imaging parameters (Groves et al., 2011). Briefly, the concatenated imaging data is decomposed into a series of linked independent components. Each imaging parameter may contribute a percentage to the variability explained by a particular linked component. Linked components are composed of a spatial map for each MRI parameter that displays where those particular metrics co-vary across participants according to a common component weighting that describes the relative variability across participants (Figure 1). Functional data could not be incorporated in our linked models because only a subset of participants’ functional data passed quality check procedures.

**Figure 1:**
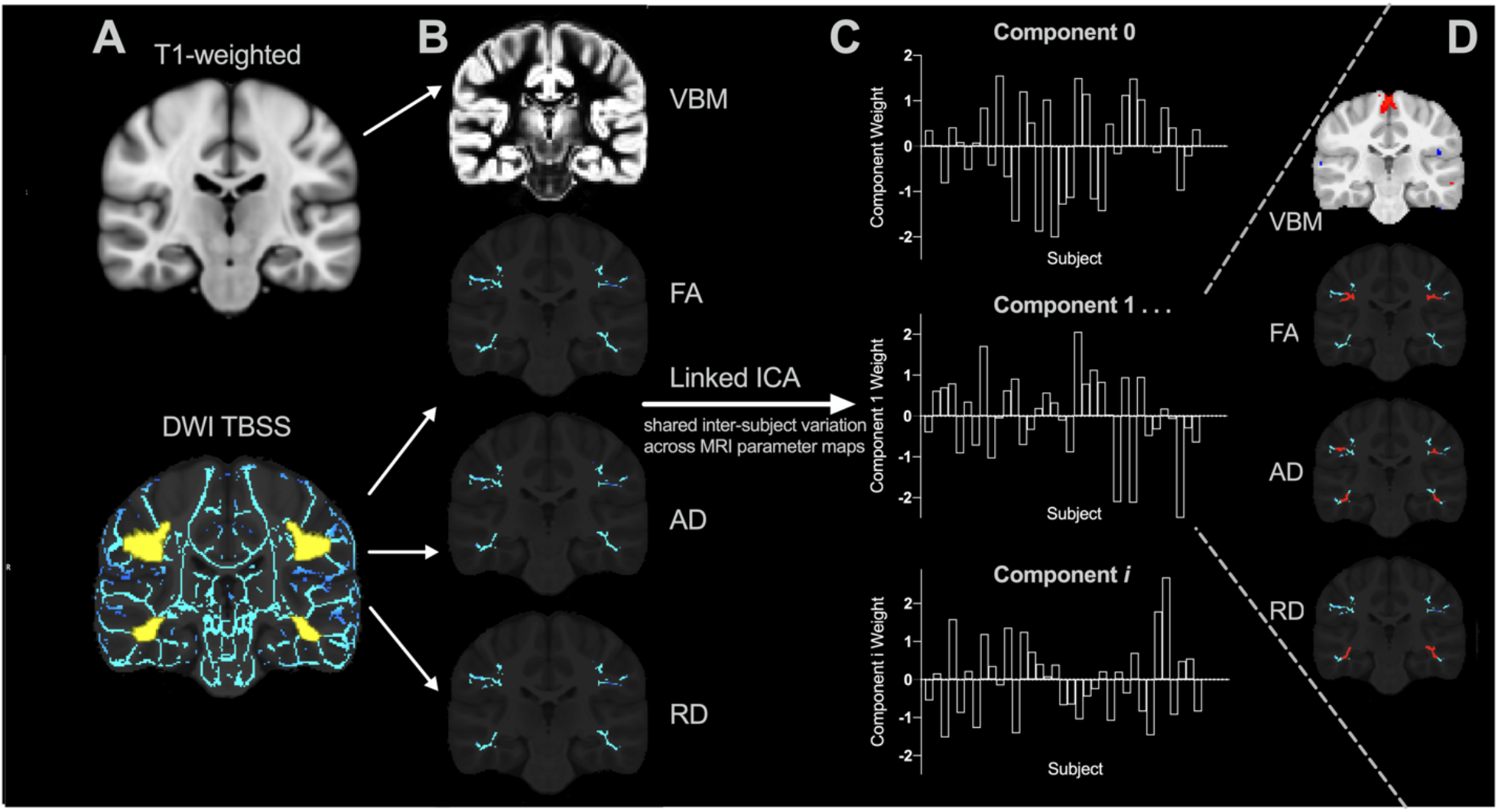
Linked independent component analysis pipeline. (A) Preprocessed T1 and diffusion weighted images were processed using VBM and DTI pipelines (with the reading-related tracts mask shown in yellow) to create (B) individual VBM, FA, AD, and RD spatial maps for each subject as input for the linked independent component analysis that decomposes the data into (C) a series of linked components comprised of component weights that reflect the inter-subject variation across (D) all input measures (VBM, FA, AD, and RD) in specific brain regions.

### Resting State Functional MRI Network Analysis

Resting state functional MRI preprocessing is described in detail elsewhere (Long et al., 2019). Briefly, processing was undertaken using FSL (slice timing, head motion correction, T1 image segmentation, head motion outlier detection, co-registration, and spatial normalization and smoothing) (Jenkinson et al., 2002) and AFNI v17.3.03 (regression of the nuisance signals, band-pass filtering and linear trend removal) (Cox, 1996). fMRI data was registered to a pediatric template in MNI standard space (Fonov et al., 2011). Participants with fMRI data that had < 4 minutes of low-motion data, or excessive motion (frame-wise displacement < 0.3mm) at any time (Satterthwaite et al., 2013) were excluded from the analysis, resulting in a total of 21 fMRI datasets.

Clean fMRI data (n = 21) was temporally concatenated and analyzed using independent component analysis to decompose the data into 20 components or networks. To identify resting state networks (RSNs), the components were compared (using cross correlation) to the 10 known RSNs (Smith et al., 2009). We identified and focused on the left-lateralized fronto-parietal language (FPL) network because it involves regions that are functionally active during cognitive-language tasks, including Broca’s and Wernicke’s areas (Smith et al., 2009; Reynolds et al., 2019b). We calculated the inter-network functional connectivity between the FPL network and the visual, default mode network (DMN) and the cerebellar RSN. Dual regression algorithms were used to back-reconstruct subject-specific RSNs and the inter-network connectivities between the FPL network and the aforementioned RSNs were calculated.

### Statistical analysis

This exploratory study utilized partial correlations, controlling for sex, and we performed bootstrapping in SPSS (version 26) with 1000 to explore the relationship and confidence interval of any relationships between the linked ICA component weights at 3.5 years and pre-reading measures at 4.5 years. Sex was included as a covariate because of known sex differences in brain development (Reynolds et al., 2019a). The inter-network functional connectivity between the FPL and the three aforementioned RSNs was also explored for correlations with pre-reading measures one year later using a partial correlation analysis controlling for sex and age at the time of the scan. As this was an exploratory study, the raw *p*-values are reported when investigating the relationships between component weights, inter-network functional connectivity and pre-reading measures.

## Results

### Language and Pre-reading Measures

At 4.5-years, the mean Phonological Processing Standard Score was 11.9 ± 2.1 and scores ranged from 7 - 16. The mean Speeded Naming Standard Score was 12.4 ± 2.3 and scores ranged from 7 - 18.

### Linked ICA

The whole-brain skeletonised DTI maps (FA, AD, RD) were input into the linked ICA with the gray matter morphometry maps. This rendered 15 components. Four components were dominated by one data set (not necessarily the same participant), and two were related to children’s age and thus were not analyzed further. Of the nine components remaining, none of the component weights were significantly related to pre-reading scores.

To be more specific to brain variabilities related to reading, we conducted a secondary analysis by masking the DTI maps to areas known to have associations with reading in older children (bilateral UF, IFO/ILF, and SLF). These were combined with the full-brain gray matter morphometry maps, which generated 14 components with the linked ICA analysis. Of the 14 components, three were dominated by one data set and two were significantly associated with age and thus were not included in the correlation analysis between the brain imaging components and pre-reading measures. Of the remaining nine components, two correlated with pre-reading measures. To confirm the validity of components of interest, linked independent component models with 13 and 15 components (model order) were also assessed. The two components described below were reproducible using all three model orders (*p* < 0.05).

Component 8 was associated with Speeded Naming (r = 0.37, *p* = 0.03, CI: [0.04,0.64]) at age 4.5 years (Figure 2), but this relationship would not survive FDR correction. This component involved volumetric variability in the angular gyrus, thalamus, fusiform gyrus, middle temporal regions, and the dorsolateral prefrontal cortex (Table 1), linked with bilateral changes in FA and AD along the SLF. Of note, higher component weights in this case represent larger volumes in the sensorimotor regions, the angular gyrus, thalamus, fusiform gyrus, and middle temporal area, smaller volumes in the posterior cingulate and visual association areas and the prefrontal cortex, and higher FA and AD with lower RD in the bilateral SLF. The component 8 weight at 3.5 years had a positive association with higher scores on the pre-reading measures at 4.5 years.

**Figure 2:**
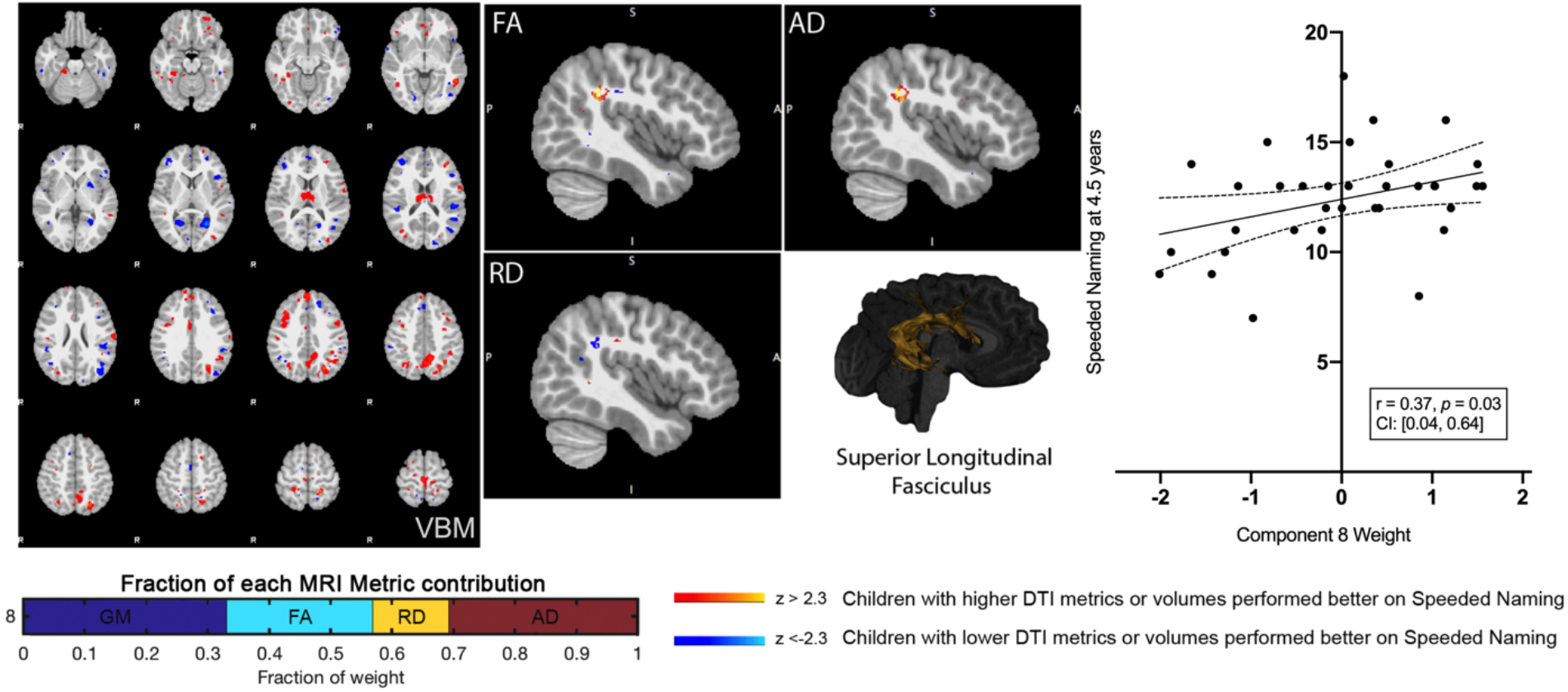
Component 8 was approximately evenly weighed by the voxel-based morphometry (VBM) gray matter, fractional anisotropy (FA), radial diffusivity (RD), and axial diffusivity (AD) contributions. This component was positively correlated with Speeded Naming at 4.5 years. For regions in red (positive clusters), children with larger volumes/higher DTI metrics performed better on Speeded Naming, and for regions in blue (negative clusters), children with smaller volumes/DTI metrics performed better on Speeded Naming. For reference, the superior longitudinal fasciculus is also shown.

**Table 1:**
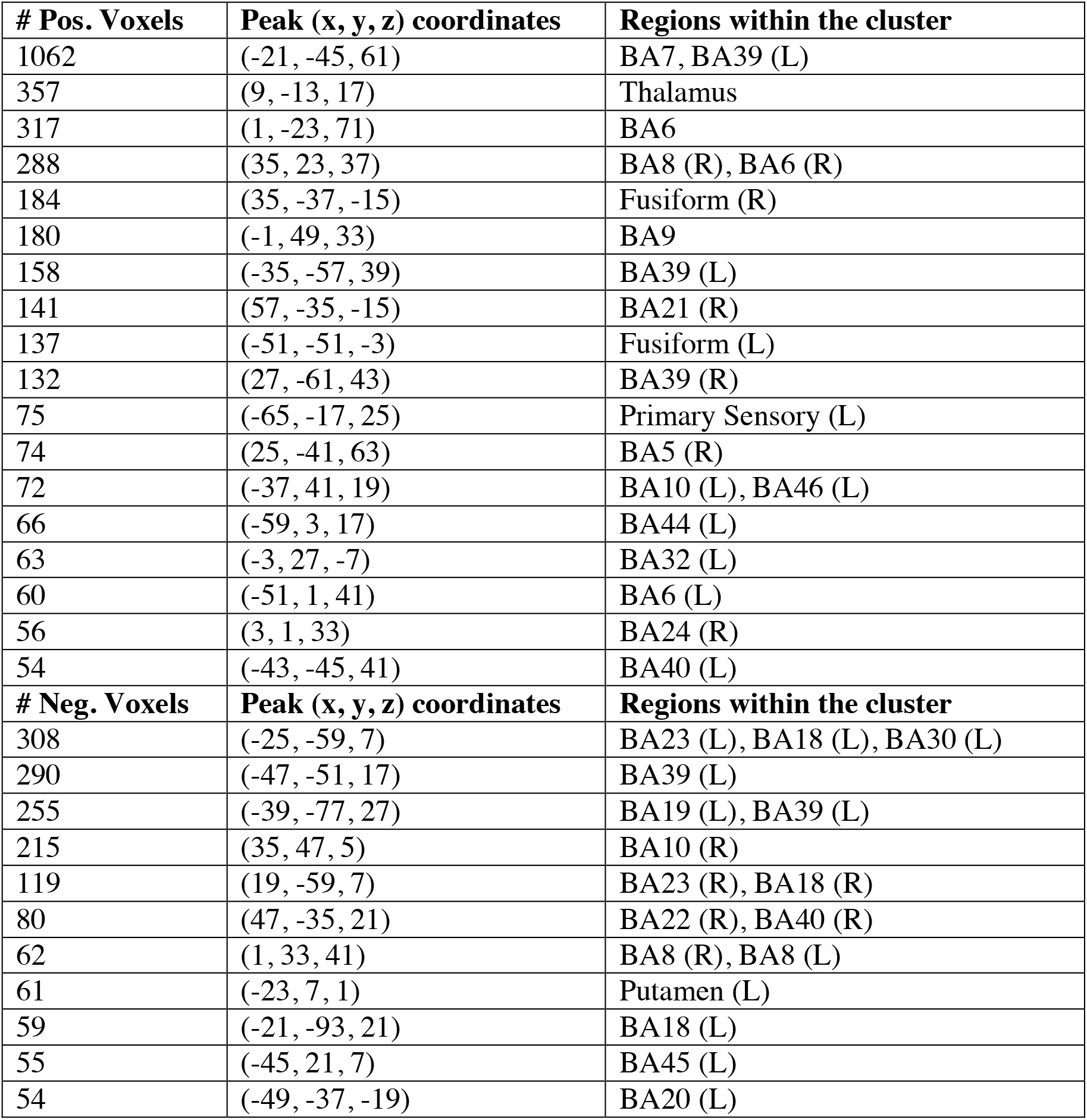
Characterization of Component 8 VBM gray matter clusters.

A second component (Component 10) was dominated by volumetric changes in the cerebellum, precuneus, angular gyri and areas in the occipital cortex (visual regions), and was associated with Speeded Naming (r = -0.35, *p* = 0.04, CI: [-0.61,-0.06]) at age 4.5 years (Figure 3) but also would not survive FDR correction. Smaller volumes in the cerebellum, and supramarginal and angular gyri, and larger volumes in visual areas in the occipital lobe and the fusiform at 3.5 years (Table 2) were associated with better pre-reading at 4.5 years.

**Figure 3:**
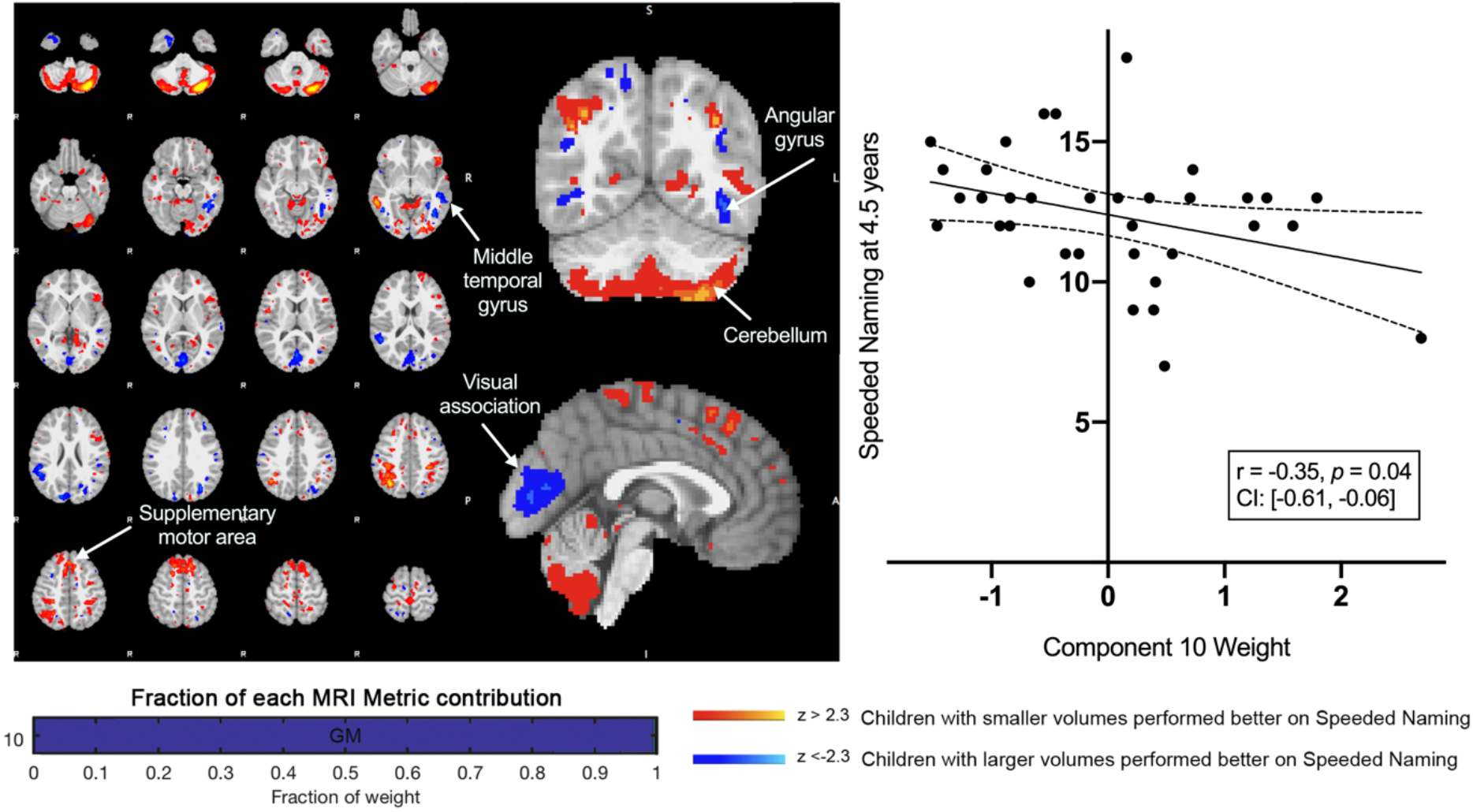
Component 10 was dominated by gray matter contributions. This component was negatively correlated with Speeded Naming performance at 4.5 years. For regions in red (positive clusters), children with smaller volumes performed better on Speeded Naming, and for regions in blue (negative clusters), children with larger volumes performed better on Speeded Naming.

**Table 2:** Characterization of Component 10 VBM gray matter clusters.

| # Pos. Voxels | Peak (x, y, z) coordinates | Brodmann Regions |
| --- | --- | --- |
| 9016 | (-29, -77, -41) | Cerebellum (L) |
| 1670 | (13, 31, 59) | BA8 (R) |
| 1186 | (35, -65, 41) | BA39 (R) |
| 669 | (-47, 33, -5) | BA47 (L) |
| 486 | (-37, -37, 45) | BA40 (L) |
| 398 | (-3, -19, 75) | BA6 (L) |
| 245 | (-15, 53, 29) | BA9 (L) |
| 234 | (59, -41, -5) | BA21 (R) |
| 167 | (-63, -3, -7) | BA22 (L) |
| 111 | (-49, -59, 7) | BA39 (L) |
| 90 | (49, 1, 13) | BA6 (R) |
| 88 | (41, -37, -19) | Fusiform (R) |
| 78 | (-13, -89, 33) | BA19 (L) |
| 76 | (15, 7, -19) | BA11 (R) |
| 73 | (19, -71, 47) | BA7 (R) |
| 69 | (-42, -6, 45) | BA6 (L) |
| 66 | (21, -37, 61) | BA5 (R) |
| 64 | (19, -9, 61) | BA6 (R) |
| 63 | (-43, -34, -12) | BA20 (L) |
| 61 | (29, 23, 41) | BA8 (R) |
| 60 | (15, -77, -5) | BA18 (R) |
| 58 | (-23, 13, -21) | BA47 (L) |
| 58 | (-49, 43, -15) | BA47 (L) |
| 55 | (29, -11, 69) | BA6 (R) |
| 51 | (21, -95, -5) | BA18 (R) |
| # Neg. Voxels | Peak (x, y, z) coordinates | Regions within the cluster |
| 1238 | (15, -87, 27) | BA19 (R) |
| 451 | (-45, -49, -15) | Fusiform (L) |
| 444 | (51, -47, 23) | BA39 (R) |
| 247 | (-29, -69, 33) | BA39 (L) |
| 243 | (-51, -31, -7) | BA21 (L) |
| 195 | (23, 5, -41) | BA36 (R) |
| 101 | (-31, -49, -5) | BA19 (L) |
| 87 | (-43, -71, 9) | BA19 (L) |
| 83 | (45, -55, -3) | Fusiform (R) |
| 67 | (15, -55, 57) | BA7 (R) |

### Resting State Networks and Pre-Reading

We examined inter-network functional connectivity between the FPL RSN (Figure 4) and three other networks including the default mode network, the occipital pole visual and the cerebellar RSNs. The inter-network functional connectivity (correlation between RSN time series) between the FPL and the DMN assessed at 3.5 years of age was correlated with Phonological Processing scores one year later at 4.5 years of age (r = 0.62, p = 0.006, CI: [0.25,0.86]) and this relationship would survive FDR correction. Also, inter-network functional connectivity between the visual RSN and the FPL at 3.5 years of age was associated with Phonological Processing scores one year later (r = 0.51, p = 0.03, CI: [0.001,0.85]).

**Figure 4:**
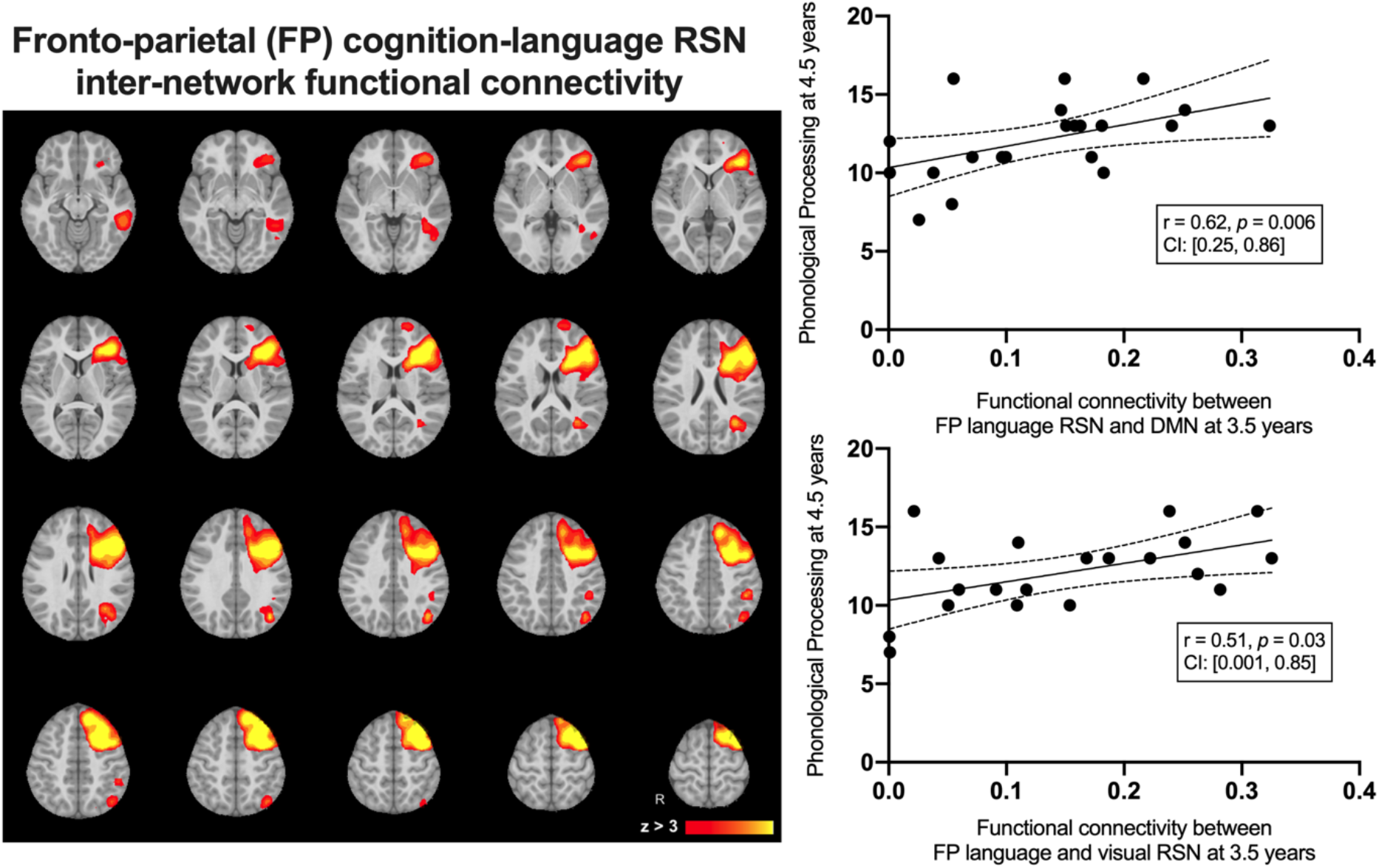
The group average left-lateralized fronto-parietal resting state network (left) and the linear relationships between inter-network functional connectivity at 3.5 years predicting Phonological Processing scores at 4.5 years of age.

## Discussion

In this exploratory study of typically developing pre-school aged children, we identified multiparametric imaging features at 3.5 years of age that were associated with pre-reading measures one year later. Our study identified variations in brain volume (angular gyrus, lingual gyrus, thalamus, fusiform gyrus, middle temporal regions, cerebellum, precuneus, and the dorsolateral prefrontal cortex), SLF microstructure, and inter-network functional connectivity of the FPL network prior to formal reading instruction at 3.5 years of age that were related to pre-reading skills at 4.5 years. Together these findings suggest that gray and white matter structure, as well as inter-network functional communication, underlie later pre-reading skills in children.

Microstructural variations (FA, AD, RD) at 3.5 years in the superior longitudinal fasciculus (SLF), together with morphological variations in cortical regions connected by the SLF, formed one imaging component that was associated with Speeded Naming scores at 4.5 years of age. Higher FA and AD, and lower RD indicate increased directionality of diffusion, and are often associated with increased myelin and/or more tightly packed axons (Beaulieu, 2002; Song et al., 2002, 2003). Because FA generally increases and RD decreases during typical early childhood development (Reynolds et al., 2019b), these profiles suggest relatively more mature white matter in the children who had better pre-reading skills at age 4.5 years.

Importantly, gray and white matter brain development are inherently linked. Our analysis allowed us to describe this co-development through the shared variability of known reading-related gray matter regions (e.g., supramarginal and angular gyrus and dorsolateral prefrontal cortex) that are structurally connected via the SLF, where we also observed coinciding variations in white matter microstructure. The SLF continues to develop into adulthood in conjunction with cerebral structural development that involves both thinning and thickening of the cortex depending on the region and stage of development (Remer et al., 2017). We observed that volumes in specific portions of Wernicke’s area and Broca’s area were linked to SLF diffusion parameter variability and together, these predicted better pre-reading scores one year later. This provides a more thorough understanding of the developing neurobiology that may support the development of pre-reading and potentially reading skills compared to single-modality studies.

Our multiparametric analysis approach demonstrated that children aged 3.5 with a relatively more mature SLF coinciding with cortical features of the angular gyrus, thalamus, fusiform gyrus, middle temporal regions, and the dorsolateral prefrontal cortex are associated with better pre-reading skills at 4.5 years of age. Though this finding would not survive FDR correction, this association between the SLF and pre-reading measures is consistent with recent findings that the microstructure of the arcuate fasciculus (a sub-section of the SLF) during infancy is associated with phonological awareness skills (Woodcock Johnson Oral Language Assessment) and vocabulary knowledge (Peabody Picture Vocabulary Test - IV) in Kindergarten (Zuk et al., 2019). The SLF, including the arcuate fasciculus, supports communication between specific regions and networks in the brain and is related to pre-reading skills. Reading-related tracts continue to develop at varying rates throughout the preschool period and the SLF for example continues to develop throughout childhood as it refines and integrates inter-network communications, which could enable more advanced reading abilities throughout childhood and adolescence (Reynolds et al., 2019b).

We also identified a second component that was associated with later pre-reading measures. This brain signature was dominated by volumetric variations in the cerebellum and the visual cortex, along with other reading-related regions within the temporal and parietal lobes. Larger visual and angular gyri cortical volumes and smaller volumes in the cerebellum, frontal and sensory regions were related to better Speeded Naming scores. The cerebellum plays a critical role in reading development through both the dorsal and ventral circuits that support phonological and semantic processes (Alvarez and Fiez, 2018; Benischek et al., 2020). Gray matter regions that were isolated in this component include regions within the dorsal and ventral cerebellum, suggesting that the structure of the cerebro-cerebellar pathway is a key developmental feature in the development of pre-reading skills. These results may suggest an earlier refinement of the cerebellum and somatosensory cortical regions combined with growth of the visual cortex that together relate to better pre-reading skills later in the preschool period at 4.5 years of age.

Phonological processing requires the use of multiple brain regions (Vigneau et al., 2006) and we found that higher inter-network functional connectivity between the FPL network and both the default mode and visual resting state networks predicted better Phonological Processing scores a year later. This suggests that a more inter-connected functional architecture lays the foundation for higher level processes that require seamless communication between a variety of networks throughout the brain. While the simple recall Speeded Naming test was associated with the microstructure of the brain’s white matter and cortex, Phonological Processing was related to the functional connectivity of the cortex. The white matter tracts that support these higher-level phonological processes are in the early stages of development during the preschool period and will continue to develop into middle childhood. As these functional brain signals are shared between specific cortical regions and networks, the underlying microstructure of pathways connecting those areas mature and refine to directly support the functional pathways. Children who utilize those pathways more often through more exposure to language and reading in early childhood may set up both the structural and functional foundations to support reading abilities later in life.

This exploratory study involved a group of typically developing children who displayed moderate to high pre-reading skills and a limited range of scores. Future studies including more children as well as those at risk for reading disorders would provide a wider range of scores and may assist in determining whether similar patterns hold in children with pre-reading difficulties that go on to display reading problems at school. Longitudinal MRI studies are a powerful way to study the developing human brain non-invasively, but require time and patience, especially with young children. Though we had a tight age range in this study, the sample size was relatively small at 35 children and the relationships were under-powered. The structural associations we found would not survive a statistical correction, so further studies with larger sample sizes are necessary to confirm these relationships between early brain measures and their influence on a child’s pre-reading abilities later in life.

## Conclusions

In this study we found that the linked structural development of white and gray matter features in early childhood were associated with pre-reading measures one year later. In particular, our analysis suggests that the structure of the SLF and cortical regions physically connected by the SLF in early childhood may play a role in the foundation for pre-reading skills. We also identified regions involved in cerebellar-cerebral reading-related circuits during early childhood that were associated with children’s performance on preschool pre-reading measures. The linked components we identified represent two developmental brain signatures that demonstrate the complex development of white matter microstructure that coincides with specific structural development of reading-related cortical regions. Phonological Processing measures were significantly predicted by inter-network communication in early childhood where we found that a more functionally integrated FPL network was predictive of better pre-reading one year later. Together, these results suggest that the co-development of white and grey matter brain structures in early life as well as the integration of functional networks before formal education are associated with pre-reading abilities in preschool children.

## Acknowledgements

This work was supported by the Canadian Institutes of Health Research (CIHR) (funding reference numbers IHD-134090, MOP-136797, New Investigator Award to C.L). C.L. receives funding from the Canada Research Chair program. K.Y.M was supported by an NSERC Postdoctoral Fellowship and the T. Chen Fong Postdoctoral Fellowship in Medical Imaging Science. J.E.R was supported by an Eyes High University of Calgary Postdoctoral Scholarship, the T. Chen Fong Postdoctoral Fellowship in Medical Imaging Science, and a CIHR Postdoctoral Fellowship (MFE-164703). AL was supported by the Horizon 2020 Programme CANDY (Grant No. 847818). The authors thank members of the APrON study for assistance with recruitment. We have no conflicts of interest to disclose.

